# Natural infection and vertical transmission of two flaviviruses (Yellow fever and Zika) in mosquitoes in primary forests in the Brazilian state of Rio de Janeiro (Diptera: Culicidae)

**DOI:** 10.1101/688713

**Authors:** Jeronimo Alencar, Cecilia Ferreira de Mello, Carlos Brisola Marcondes, Anthony Érico Guimarães, Helena Keiko Toma, Amanda Queiroz Bastos, Shayenne Olsson Freitas Silva, Sergio Lisboa Machado

## Abstract

**Background:** Zika virus (ZIKV) was recently introduced in the American continent, probably transmitted by *Aedes aegypti* and possibly by *Ae. albopictus* and *Culex quinquefasciatus* in urban environments. ZIKV represents a known public health problem as it has been involved in newborn cases of congenital microcephaly in South America since 2005. The transmission of this virus in forested areas of other countries and its relative ubiquity in relation to its vectors and reservoirs raises suspicions of its adaptation to non-human modified environments (*i.e*., natural forests reserve) or on this continent, similar to those seen for Yellow fever virus (YFV). The objective of this work was to have an epidemiological monitoring tool mapping insects as well as circulating arboviruses in wild areas with low human interference. This study was based on the history of the insect flavivirus spreading cycle.

**Methods/Principal Findings:** Using a previously described sensitive PCR-based assay to assess the conserved NS5 region of the *Flavivirus* genus, both YFV partial genome and ZIKV were found in pools of *Aedes albopictus*, a sylvatic mosquito adapted to human-modified environments, and in *Haemagogus leucocelaenus*, a sylvatic mosquito.

**Conclusions:** This is the first report of natural infection by ZIKV in mosquitoes in a sylvatic environment on the American continent. The wide distribution of these mosquitoes is probably important in the transmission of ZIKV. Vertical transmission indicates a higher efficiency for the maintenance and transmission of the virus in nature as well as the presence of the ZIKV in permanent character in the forest areas as it occurs with the YFV thus making more difficult the prevention of new cases of Zika in humans.

**Author Summary:** Arboviruses are diseases transmitted by arthropod vectors, hence the origin of the term ARthropod BOrne VIRUS, which is adopted since 1942. This work had as objective to survey the circulating insects as well as to detect the presence of viruses in them. Arboviruses circulate between insects and vertebrate hosts, having importance for promoting diseases in humans and animals. The diseases most known at the time, due to the recent cases reported by South America, are Dengue, Zika, Yellow Fever and Chikungunya. For this study, we used appropriate traps to collect the insects and their eggs in wild areas where there is little human interference. After collection, mosquitoes and / or eggs were identified and separated as to the source and species. The eggs were kept in laboratory conditions for the hatching of new insects. All the insects obtained were separated into pools to be macerated and thus extract the RNA from the viruses to be studied. Using molecular biology techniques, in our case the RT-PCR (Reverse Transcriptase Polymerase Chain Reaction), we amplified the RNA and in sequentially, we performed the sequencing reaction. With sequencing, it is possible to identify which virus material is present since each virus has a characteristic arrangement. For the identification of the sequences, we need to use some computational programs that guarantee us the correct result.

## Introduction

The Flaviviridae family has four genera and the *Flavivirus* genus contains over 90 viruses, some of which are of clinical importance for humans and animals, such as the Yellow fever virus (YFV), Dengue virus (DENV), Zika virus (ZIKV), and West Nile virus (WNV). Flaviviruses are enveloped, icosahedral viruses and have an RNA genome composed of a single positive-strand chain of approximately 11000 nucleotides. Their genomes encode three structural proteins (capsid [C], envelope [E], and pre-Membrane/membrane [prM/M]) and seven non-structural proteins (NS1, NS2a, NS2b, NS3, NS4a, NS4b, and NS5) that have functions involved in replication, virulence, and pathogenicity [1].

ZIKV, first identified in a forest in Uganda, has recently spread to Asia, Pacific islands, and the American continent, causing serious concern for microcephaly in fetuses [2–4]. Mostly transmitted in urban environments by *Aedes aegypti* and possibly *Aedes albopictus*, it has been found in many species of mosquitoes, mostly in *Aedes* species (31 spp.), but also in some *Anopheles, Culex* and other genera, and even in a species of horse fly [2,5]. The first autochtonous case of ZIKV in Brazil was diagnosed in May 2015[6], and its circulation was confirmed in all 26 states and federal district [7].

Molecular techniques, as well as immunological methods, have been used to diagnose ZIKV in a variety of species of mammals belonging to nine orders (Artiodactila, Carnivora, Cetartiodactyla, Chiroptera, Lagomorpha, Perissodactyla, Primates, Proboscidea and Rodentia), three orders of birds (Anseriformes, Charadriiformes, and Ciconiiformes) and lizards Squamata [8], mostly in Africa. On the American continent, possibly due to its recent introduction and smaller number of studies, it has only been found in monkeys in Ceará state [9]. Some primates and Edentata have been found to be serologically positive for *Flavivirus* in the South of Bahia state [10].

Even though non-human primates (NHPs) are considered of low importance, their circulation in low degradation forest environments, especially the diversified fauna of mosquitoes and mammals found in Brazil, needs to be assessed [8, 11].

In this study, we report the natural infection and vertical transmission of ZIKV and YFV in *Haemagogus leucocelaenus* and *Ae. albopictus* in a forest in Casimiro de Abreu and Nova Iguaçú, in the Brazilian state of Rio de Janeiro.

There is currently an ongoing outbreak of sylvatic yellow fever in Brazil. The outbreak probably started at the end of 2016, when the first case was reported in the state of Minas Gerais, but has since spread to the states of Espírito Santo, São Paulo, and Rio de Janeiro. According to a WHO report, as of April 2017, YFV transmission (epizootic and human cases) continues to expand towards the Atlantic coast of Brazil in areas not previously deemed to be at risk for yellow fever transmission [12].

The main genera of mosquitoes capable of being infected and transmitting the sylvatic YFV are *Haemagogus* and *Sabethes* which are considered to be biological vectors and to be responsible for maintenance of the natural cycle of this zoonosis in forested areas of the Americas. In the southeast of Brazil, during the present epidemic, *Hg. leucocelaenus* and *Hg. janthinomys* are considered important vectors [13].

*Haemagogus* are essentially sylvatic mosquitoes, with diurnal acrodendrophilic habitats, and are mainly found in densely forested areas [14]*. Haemagogus leucocelaenus* is the species most frequently found in Brazil and is considered a primary vector for SYF in southeastern Brazil. It is widely distributed from Trinidad to the south of Brazil, This Culicidae is commonly found in Brazil and is considered to be of epidemiological importance due to its involvement in the transmission of arboviruses, with Yellow fever being one of the most important. A group of researches [15] reported through the use of hemi-nested reverse transcriptase PCR identifying sequences compatible with DENV-1 in *Hg. leucocelaenus* from Coribe, a city from northeast Bahia, suggesting the occurrence of a sylvatic cycle and highlighting the importance of studies regarding these wild mosquitoes [16,17].

## Materials and methods

### Ethics statement

All research was performed in accordance with scientific license number 44333 provided by (Ministry of Environment - MMA, Chico Mendes Institute of Biodiversity Conservation - ICMBio, Biodiversity Information and Authorization System – SISBIO). Forests collections were undertaken with informed consent and cooperation of the property owners, householders or local authorities. All members of the collection team were adequately vaccinated against YFV and were aware of the potential risks in the area under study.

### Study areas

The eggs of *Hg. leucocelaenus* and *Ae. albopictus* were collected using ovitraps, from September 2018 to March 2019. The present study was carried out in Três Montes Farm (TMF), Três Morros Private Reserve of Natural Heritage - TMPRNH, Casimiro de Abreu city), and Sítio Boa Esperança (Tinguá, Nova Iguaçu city). Casimiro de Abreu located in southeastern Brazil, approximately 140 km from the city of Rio de Janeiro and Tinguá at approximately 30 km from Rio de Janeiro. Samples were collected from four sampling sites. Geographical coordinates of the sampling sites were obtained using a Garmin GPSMAP 60CS (Garmin International, Inc., Olathe, KA, USA) (Table 1).

**Table 1.** Detection of Yellow fever virus (YFV), Zika virus (ZIKV) in *Aedes albopictus* and *Haemagogus leucocelaenus* in primary forests in the Brazilian state of Rio de Janeiro, Brazil.

| Pool | Mosquito Species | Sex | Total specimens | Month/Year collected | Geographic coordinates | Trap identification | PAN Flavivirus PCR Result | Sequencing result |
| --- | --- | --- | --- | --- | --- | --- | --- | --- |
| 43 | *Ae. albopictus* | ♀ | 32 | jan/19 | 22°33'01.3" S<br>42°00'52.7" W | TMPR NH - 32* | Positive | Zika Virus |
| 45 | *Ae. albopictus* | ♂ | 2 | Oct/2018 | 22°31'40.1" S<br>42°02'58.6" W | TMF -2 * | Positive | Zika Virus |
| 54 | *Hg. leucocelaenus* | ♀ | 6 | jan/19 | 22°35'11.98" S<br>43°24'34.12" W | Tinguá* | Positive | Zika Virus |
| 62 | *Ae. albopictus* | ♀ | 9 | Oct/2018 | 22°31'43.9" S<br>42°02'56.8" W | TMF -3 * | Positive | Yellow Fever |
| 64 | *Hg. leucocelaenus* | ♂ | 3 | Sep/2018 | 22°31'49.5" S<br>42°02'56.3" W | TMF -7 * | Positive | Yellow Fever |
|  |  |  | 5 | Oct/2018 |  |  | Positive | Yellow Fever |

\* (Três Montes Farm – TMF, sites 2, 3, and 7), \* (Três Morros Private Reserve of Natural Heritage– TMPR NH, site 32), and \* (Sitio Boa Esperança, Tinguá, Nova Iguaçu).

Location of the study area and sampling sites in Rio de Janeiro. Geographical coordinates of the sampling sites were obtained using the Garmin GPSmap 60 CS GPS. Maps were prepared in ArcGIS PRO data in the public domain (URL: https://pro.arcgis.com/en/pro-app/. Accessed: May 2019) and edited in CorelDRAW Graphics Suite X7 (Fig 1).

**Fig 1.**
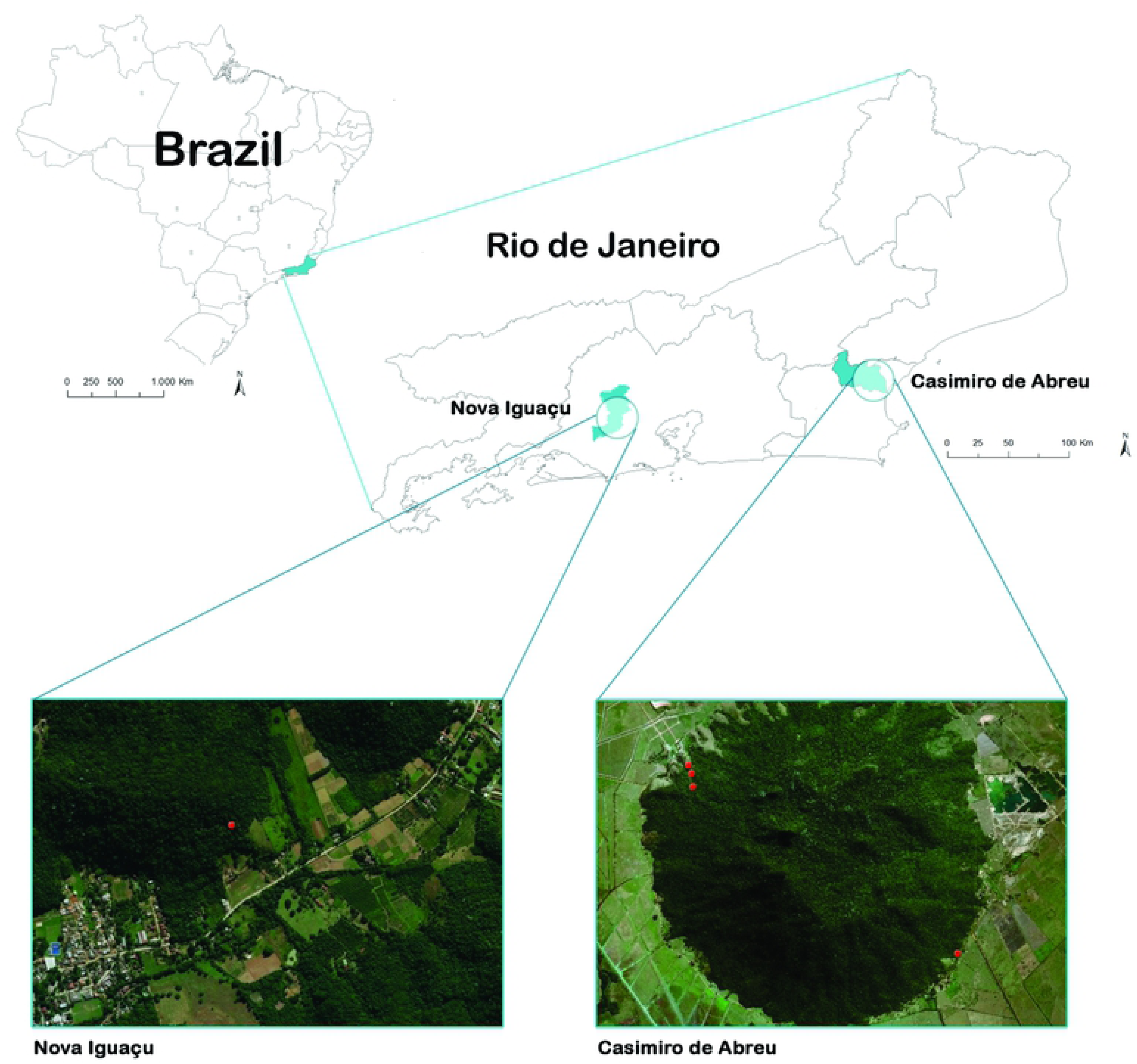
Map of the locations used for mosquito collection in the study. (Red: Positive sampling sites for Yellow fever (YFV) viruses, Zika virus (ZIKV) in primary forests in the Brazilian, municipalities of Nova Iguaçu and Casimiro de Abreu, state of Rio de Janeiro, Brazil). Maps were prepared in ArcGIS PRO data in the public domain (URL: https://pro.arcgis.com/en/pro-app/. Accessed: May20I9) and edited in CoreIDRAW Graphics Suite X7.

The main land cover of the region is typical Atlantic forest vegetation, with dense ombrophilous sub-mountain forests in moderate and advanced stages of regeneration. The region, located in the hydrographic basin of São João River, is situated in the intertropical zone (at low latitudes) and is highly influenced by the Atlantic Ocean. Thus, its climate is predominantly a humid tropical type [18]. The average temperature is 26.8°C, with a relative humidity of 56% and 1,200 mm precipitation [19]. Higher rainfall levels occur from October to March.

At each sampling site, samples were collected using ovitraps. The ovitraps consisted of cylindrical 1L black plastic containers, containing four wooden paddles (2.5 × 14 cm). The traps were placed at a height that varied between 2 and 10 m above soil level. Details on the use and manufacturing of the ovitraps can be found in studies previously done [20,21]. The paddles in the traps were examined every two weeks to detect and quantify the eggs.

Just after arriving in the laboratory positive-paddles were immersed in white trays filled with dechlorinated water at 29 ± 1°C and these trays were kept in an acclimatized chamber for hatching. After 3 days, the paddles were removed from the water and left to air dry for another 3 days to quantify the hatched larvae. Immature forms were reared as previously described [22], and processed for diagnosis of natural infection by arboviruses through molecular techniques.

Before extraction, all mosquitoes were retained in cryotubes in a −80°C ultrafreezer, separated by species and trap identification. The specimens were kept alive for specific determination in adulthood, by direct observation of the morphological characters evidenced by the stereomicroscopic microscope (Zeiss®) and consultation with the respective descriptions/ diagnoses of the spp, using dichotomous keys [14, 23]. Mosquito genera were abbreviated according to a well-established abbreviation [24].

### RNA extraction

We made pools of 3 to 33 mosquitoes, mixing males and females to be further macerated. Viral RNA was extracted from the mosquito pools using MN Nucleo Spin® RNA (Macherey-Nagel GmbH & Co. KG, ref. 740955.250) and cDNA was immediately synthetized using Hi-capacity RNA-to-DNA™ kit (Applied Biosystems™, ref. 4388950) both according to manufacturers instructions. DNA was then quantified using Denovix DS-11+b Quantifier (DeNovix Inc., USA) and maintained at −20°C until molecular investigation of the flaviviruses.

### PCR for *Flaviviruses*

We used primer sequences derived from the conserved NS5 region of the *Flavivirus* genus that had previously been described [25], and then adapted the conditions for PCR amplification.

The conditions used were 1 x PCR buffer, 1.5 mM MgCl_2_, 10 pmol Pan-Flavi Forward primer (5′-TAC AAC ATG ATG GGG AAR AGA GAR AA-3′) and 10 pmol Pan-Flavi Reverse primer (5′-GCW GAT GAC ACM GCN GGC TGG GAC AC-3′), 1.0 U of DNA polymerase (Thermo Fisher Scientific, 168 Third Avenue, Walthan, MA 02451, United States), and a final extension of 72°C for 5 min.

PCR products were evaluated by electrophoresis on 1.5% agarose gels in 1 × TBE (Trizma, boric acid, EDTA) buffer and visualized under UV light (260 nm) after ethidium bromide staining. The expected amplified fragment ranged from 200-300 bp and were purified using Cellco PCR purification kit (Cellco Biotec do Brasil Ltda. Cat.#DPK-106L).

### Nucleotide sequencing

Sequencing was kindly performed by the Oswaldo Cruz Foundation (FIOCRUZ) at the RPT01A sequencing laboratory as previously described [26]. Approximately 10– 40 ng of purified PCR product was sequenced following the BigDye Terminator v.3.1 Cycle Sequencing protocol using an ABI 3730 DNA Sequencer. The sequences were then analyzed using Geneious R10 (Biomatters, v.10.2.6). The contigs were compared with reference sequences using the nucleotide Basic Local Alignment Search Tool (BLASTn, GenBank, PubMed).

**Accessions** – Nucleotide sequences from the NS5 segments obtained in the present study were deposited in GenBank.

## Results

Since September 2018, 924 insects were collected and stored in a −80°C ultrafreezer until processing. Insects were separated based on the ovitrap location and species. The number of mosquitos in each pool processed was based on previous literature reports [27–30] as well as the efficiency of RNA extraction kit; as a result, we used mosquito pools raging from 3 to 33 insects. The processing of mosquito pools requires a sensitive molecular method, since the minimum infection rate in females mosquitoes ranges from 0.1 to 3.9 per 1000 [31]. Up until now, we have processed 70 pools from Três Montes Farm (TMF), Três Morros Private Reserve of Natural Heritage - TMPRNH, and Sítio Boa Esperança.

We were able to identify five positive mosquito pools from different species by carrying out the PCR using the NS5 region primers previously described [25]. When the PCR products were sequenced and analyzed by NCBI Blast (Basic Local Alignment Search Tool, at https://blast.ncbi.nlm.nih.gov/Blast.cgi), three pools positive for YFV and two pools positive for ZIKV were found. Although our PCR product sequences varied from between 200 and 241 bp, we found the sequenced pools had scores higher than 200 and identity indexes of between 90 to 96.4% of identity covering 94 to 98% of the sequences (Table 1), Gene Bank accession numbers: MK972825, MK972826, MK972827, MK972828 and MK972829.

## Discussion

The NS5 region of the Flaviviridae family was used in this study because it is a commonly conserved region and has previously been used to identify viruses using sequencing methods [25,32–36]. Through this sequencing approach, we identified ZIKV and YFV in different sylvatic mosquito species. Sylvatic Yellow fever is usually found in *Ae. albopictus, Haemagogus leucocelaenus,* and *Hg. janthinomys* species [37–38]. It is also important to understand that the complete sylvatic cycle can be maintained in forests in the presence of vertebrate reservoirs [39]. A recent publication showed that *Aedes* mosquitoes can also be contaminated with ZIKV by breeding in contaminated aquatic environments [40].

Here, for the first time, ZIKV has been found in a forest environment on the American continent in both the sylvatic mosquito *Hg. leucocelaenus* and *Aedes albopictus*, a mosquito adapted to sylvatic and urban environments. It has been previously reported that *Aedes albopictus* is a natural ZIKV vector in several countries [41–44].

The finding of *Hg. leucocelaenus*, a sylvatic species, infected with ZIKV indicates the circulation of virus in this area, presumably along with some vertebrate reservoir(s). *Aedes albopictus* presents a transmission potential for ZIKV [45] and the simultaneous finding of this mosquito in the area means a risk for spillover from the forest to human-modified environments [46].

The blood feeding sources of *Hg. leucocelaenus* are diverse, feeding mostly on birds, but also on several mammals in Rio de Janeiro and Goiás state [22]. *Aedes albopictus* feeds mostly on mammals, preferring human blood when available [47, 48], but is also ubiquitous in its absence [49, 50].

Another risk factor is that *Ae. albopictus* can be the vector, of at least, for two different Flaviviruses since it was found in insect pools from Africa. In addition, through laboratory tests of co-infection and super-infection, the possibility of dual infection transmission to humans has been shown [51].

The present results indicate a sylvatic cycle for Zika virus; hence, it is important to investigate its presence in other vectors, especially near urban areas. Since *Ae. albopictus* can easily go from forest environments to peridomestic areas and vice versa in Rio de Janeiro [52], there is a high likelihood of viral transport between these areas.

Localities in Ceará and Bahia, where ZIKV has been found in mammals, should have their mosquito fauna carefully studied. It should be emphasized that the presence of *Hg. janthinomys*, the other highly suspected species [13] in an urban forest (Parque Dois Irmãos) in Recife, Pernambuco state [53], a city highly endemic to ZIKV and other viruses (DENV and CHIKV) [54] should be studied for natural infection.

The finding of infection in mosquitoes reared from eggs obtained under natural conditions indicates the occurrence of transovarial transmission of ZIKV, and the presence of virus in the salivary glands of these mosquitoes should be investigated. Vertical transmission has been found in many arboviruses [55], and its role in maintenance of these viruses in nature should be evaluated. A low magnitude and long duration viremia (the ‘tortoise’ strategy), has been shown to result in higher rate of persistence in vectors and host populations compared to high magnitude, short duration viremia (the ‘hare’ strategy) [56].

The vectorial competence of these species and others present in the area needs to be carefully evaluated, to elucidate the real role in the transmission of ZIKV in low degradation environments with low human presence, like forest reserves and the possibility of transference between these and modified areas. The competence of many species, mostly in *Aedes* and *Culex*, has already been tested, but not *Haemagogus* [57].

A previous study utilizing CDC traps in a reserve at Casimiro de Abreu indicated the presence of at least 15 species, not including *Hg. leucocelaeunus* and *Ae. albopictus* [58], probably due to their predominantly diurnal activity. Among the known vectors of sylvatic yellow fever, the species found in Casimiro de Abreu were *Hg. leucocelaenus* and *Hg. janthinomys*, being that this first taxon was found to be naturally infected by YFV in the same place and time as the outbreak of the disease. The results confirm the role of *Hg. leucocelaenus* as an important YFV vector in Southeastern Brazil.

Alencar et al (2016) [21] have reported three epidemiologically important mosquito species in the transmission of arboviruses (*Hg. leucocelaenus, Hg. janthinomys*, and *Ae. albopictus*) in this present study region. Considering that each viremic monkey can infect hundreds of mosquitoes, it is crucial to understand the dynamics of transmission. It is possible that a large number of *Hg. leucocelaenus* are infected and through vertical transmission, could play a role in the maintenance of epizootic and human infection, which contributes to the spread of the virus to other areas [59].

In general, YFV transmission occurs within forests, mainly affecting humans involved in activities such as logging, fishing, hunting and so on, but in the case of *Hg. leucocelaenus*, which tends to leave the forest, it can infect humans of both sexes and various ages. *Haemagogus leucocelaenus* infected with YFV were captured at ground level, where most of our specimens were obtained, during the outbreak that occurred in Rio Grande do Sul (Brazil) between 2008 and 2009 [60], which reinforces our hypothesis.

The occurrence of yellow fever virus in natural conditions demonstrates its current circulation in the Atlantic forest areas of the municipality of Casimiro de Abreu, Rio de Janeiro state. This is the first report of the detection of yellow fever virus in *Hg. leucocelaenus* in Rio de Janeiro state since the last report 88 years ago in urban YF in this state [61].

Evidence of active sylvatic SYFV transmission in the nature reserves studied here and the abundance of the main mosquito vector for this disease in Brazil, necessitates active surveillance for the emergence of this virus in neighboring communities. Forests near human-modified areas positive for arbovirus, such as urban forests (e.g., Tijuca - Rio de Janeiro; Buraquinho – João Pessoa; Dois Irmãos – Recife) are a priority.

Mosquitoes adapted to urban environments, mostly *Ae. aegypti*, transmit YFV and ZIKV among humans. Since both are well-adapted to mosquitoes of several species, the spillover to preserved forests, circulating among wild vertebrate reservoirs and mosquitoes should not be surprising. However, if not studied, such sylvatic cycles will probably be uncovered Low levels of reactivity of primates infected with ZIKY or YFV near urban areas [11] must not discourage additional studies in such areas.

These results corroborate the warning of [56] of adaptation of ZIKV to forest environments, making impossible the eradication of virus from the continent and reinforcing the need for the control of urban mosquitoes and the development of a good vaccine.

## Acknowledgements

The authors thank Farms Reunidas Agropecuarias Três Montes; Farm Três Morros, for providing the facilities to conduct the present study.

